# Physiological and transcriptomic analysis of yellow leaf coloration in *Populus deltoides* Marsh

**DOI:** 10.1101/463224

**Authors:** Shuzhen Zhang, Xiaolu Wu, Jie Cui, Fan Zhang, Xueqin Wan, Qinglin Liu, Yu Zhong, Tiantian Lin

## Abstract

As important deciduous tree, *Populus deltoides* Marsh possesses a high ornamental value for its leaves remaining yellow during the non-dormant period. However, little is known about the regulatory mechanism of leaf coloration in *Populus deltoides* Marsh. Thus, we analyzed physiological and transcriptional differences of yellow leaves (mutant) and green leaves (wild-type) of *Populus deltoides* Marsh. Physiological experiments showed that the contents of chlorophyll (Chl) and carotenoid are lower in mutant, the flavonoid content is not differed significantly between mutant and wild-type. Transcriptomic sequencing was further used to identify 153 differentially expressed genes (DEGs). Functional classifications based on Gene Ontology enrichment and Genomes enrichment analysis indicated that the DEGs were involved in Chl biosynthesis and flavonoid biosynthesis pathway. Among these, geranylgeranyl diphosphate (CHLP) genes associated with Chl biosynthesis showed down-regulation, while chlorophyllase (CLH) genes associated with Chl degradation were up-regulated in yellow leaves. The expression levels of these genes were further confirmed using quantitative real-time PCR (RT-qPCR). Furthermore, the measurement of the main precursors of Chl confirmed that CHLP is vital enzymes for the yellow leaf color phenotype. Consequently, the formation of yellow leaf color is due to disruption of Chl synthesis and catabolism rather than flavonoid content. These results contribute to our understanding of mechanisms and regulation of leaf color variation in poplar at the transcriptional level.

## 1. Introduction

Leaf color is an important feature of ornamental plants, and trees with colored leaves have been widely cultivated in landscape gardens. The main factors that determine foliage color are the types of pigment and their relative concentrations. The formation of red leaves is the result of anthocyanin accumulation, which has been extensively studied [1]. In contrast, there are only a few studies focus on the mechanism of yellow leaves. Leaf yellowing is generally considered to be caused by decreased Chl content, since Chl is the main pigment content of green leaves [2]. Therefore, studies of leaf yellowing have mostly focused on Chl biosynthesis and degradation. In addition, leaf yellowing may be also due to the accumulation of flavonoids such as flavanol, flavonol, chalcone, aurone [3,4].

The Chl biosynthetic pathway consists of about 20 different enzymatic steps, starting from glutamyl-tRNA to Chl a and Chl b [5]. Mutations in any one of the genes of the pathway can affect the accumulation of Chl [6], decrease photosynthesis capacity [7] and affect the development of chloroplast [8]. The silence of *CHLD* and *CHLI* (magnesium chelatase subunit D and I) induced by virus in peas resulted in yellow leaf phenotypes with rapid reduction of photosynthetic proteins, undeveloped thylakoid membranes, altered chloroplast nucleoid structure and malformed antenna complexes [9]. Moreover, in rice, *PGL_10_* encoded protochlorophyllide oxidoreductase B (PORB), pale-green leaf mutant pgl_10_ had decreased Chl (a and b), carotenoid contents, as well as grana lamellae of chloroplasts compared with the wild-type [10]. In addition, mutants with disrupted Chl degradation were used to characterize many steps in the Chl degradation pathway in leaves undergoing senescence [11]. In Arabidopsis mutant deficient in PPH (pheophytinase), Chl degradation is inhibited, and the plants exhibit a typeC stay-green phenotype during senescence [12]. Previous studies revealed that chlorophyllase (Chlase) is involved in Chl degradation in ethylene-treated citrus fruit and could regulate the balance between different plant defense pathways, enhance plant resistance to bacteria [13-15]. Recently, Mutants deficient in Chl biosynthesis and degradation have been identified in many yellow leaf plants, such as rice [16-19], Arabidopsis thaliana [20] and pak-choi [21].

The genotype we reported is a kind of *Populus deltoides* Marsh (mutant), which is a bud mutation of green leaf *Populus deltoides* Marsh (wild-type) (Figure 1). The mutant is a rare yellow leaf variety which was found in poplar plants of the Salicaceae family. There exists extremely high ornamental value for this species because its leaves remain golden in spring, summer and autumn. However, the molecular mechanism underlying the leaf color of the mutant has not yet been elucidated. Many ornamental plant cultivars with fruit or flower color variation arose from the bud mutation. For instance, the color in grape skin changes from white to red due to bud mutant [22], flower color mutants of roses, carnations and chrysanthemums have also been reported [23]. In contrast, yellow leaf phenotype caused by bud mutant were hardly reported. On the other hand, the study related to yellow leaf color were mostly focused on leaf yellowing. For example, the tea cultivar ‘Anji Baicha’ produces yellow or white shoots at low temperatures, and turn green when the environmental temperatures increase [24]. Only a few studies have reported the yellow leaf phenotype, such as the cucumber Chl-deficient golden leaf mutation [25].

**Figure 1.**
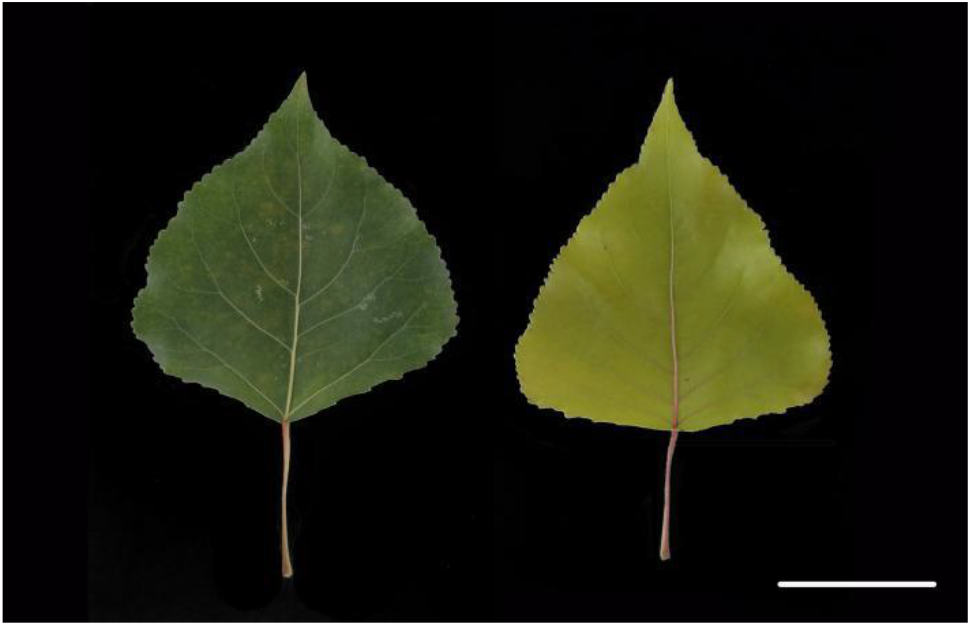
Phenotype of green leaf and yellow leaf. Bar=5 cm.

In this study, the photosynthetic pigments contents, Chl precursors contents, flavonoid contents and transcriptomics of the mutant type and wild type were analyzed. Based on a combination of biochemical analysis and bioinformatics, we identified differentially expressed genes (DEGs related to Chl and flavonoid biosynthesis. Furthermore, the expression of DEGs involved in leaf coloration was validated using quantitative real-time polymerase chain reaction (RT-qPCR). Our results clarified the physiological and transcriptomic aspects of golden leaf coloration in *Populus deltoides* Marsh and will serve as a platform to advance the understanding of the regulatory mechanisms underlying the leaf color formation in poplar and other plant species.

## 2. Results

### 2.1. Pigment content analysis of wild-type and mutant

We analyzed changes in the pigment contents of wild-type leaves and mutant leaves. The uroporphyrinogen III (Urogen III) content of the yellow leaves was significantly higher than that of the wild-type, whereas there were no significant differences in coproporphyrinogen III (Coprogen III) (Figure 2). Further detailed analysis showed that the protoporphyrin IX (Proto IX), magnesium protoporphyrin IX (Mg-Proto IX) and protochlorophyllide (Pchlide) contents of the mutant were significantly decreased by about 52.53%-64.71% than those from green leaves. On the other hand, the content of chlorophyllide (Chlide) a in yellow leaves was lower than that of green leaves. Compared with the green leaves, the Chl a content, Chl b content and carotenoids content of yellow leaves decreased significantly by 72.41%, 84.86% and 53.88%, respectively (Figure 2). In addition, the difference between the total flavonoid contents of the green leaves and yellow leaves was not significant (Figure 2).

**Figure 2.**
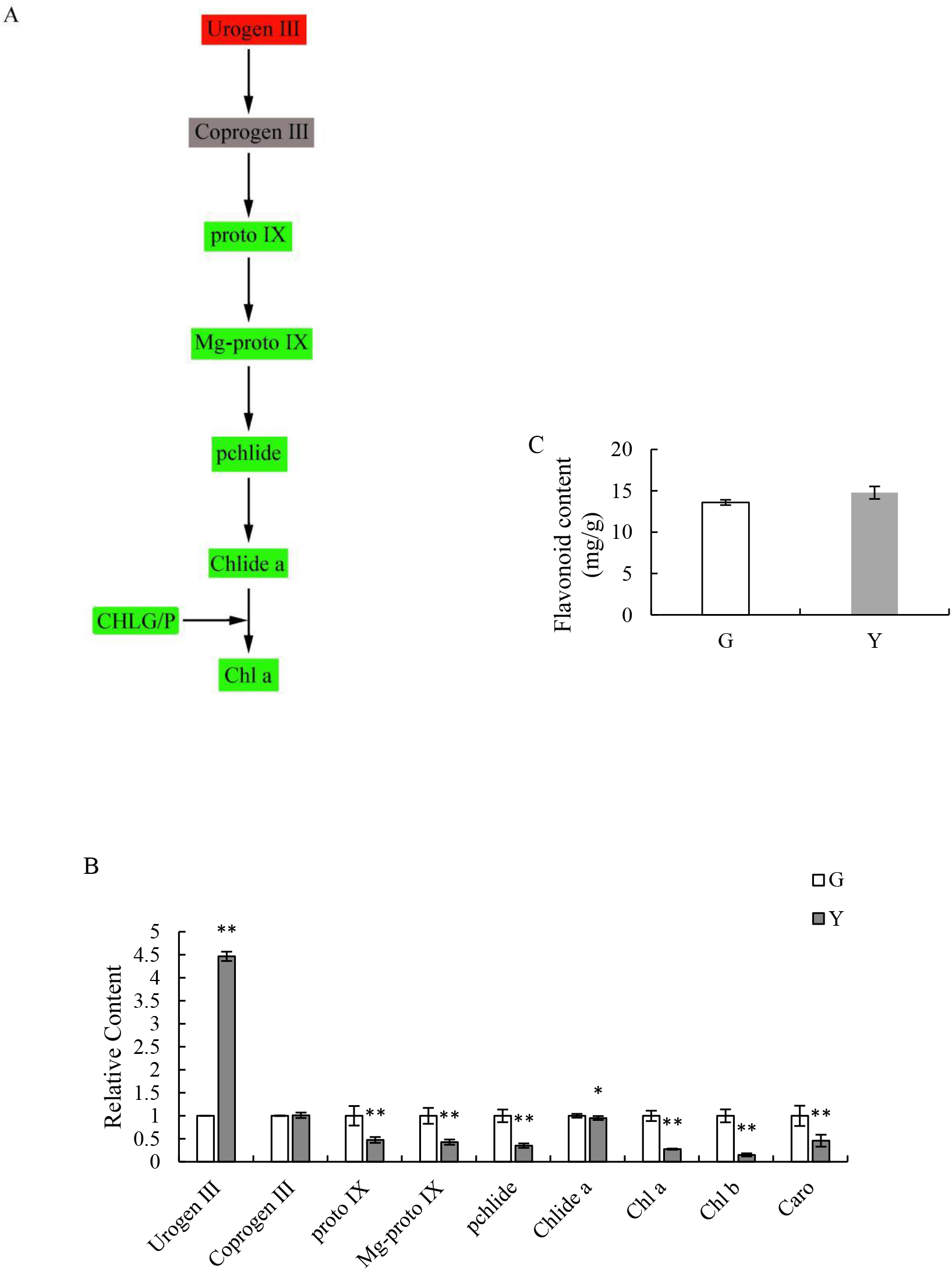
Determination of pigment contents in G (green leaves) and Y (yellow leaves). (A) Schematic view of the Chl biosynthesis pathway. The rounded rectangle shows the gene encoding protein catalyzing the reaction of the precursors. The red color means significantly increased in the Y leaves. The green color means significantly decreased or down-regulated in the Y leaves. The gray color means there was no significant difference between the G and Y leaves; (B) Comparison of the relative contents of Chl precursors and photosynthetic pigments; (C) Comparison of the flavonoid contents. Asterisks indicate: (*) P ⩽ 0.05, (**) P ⩽ 0.01. Urogen III, uroporphyrinogen III; Coprogen III, coproporphyrinogen III; Proto IX, protoporphyrin IX; Mg-Proto IX, Mg-protoporphyrin IX; Pchlide, protochlorophyllide; Chlide a, chlorophyllide a; Chl a, chlorophyll a; Chl b, chlorophyll b; Caro,carotenoid.

### 2.2. Analysis of sequencing data

RNA-seq libraries were constructed from green and yellow leaf samples and sequenced using the Illumina HiseqTM 4000 platform for acquiring a comprehensive overview of leaf coloration. Approximately 45 million and 47 million raw reads were obtained from each sample. After removal of adaptor sequence and low quality reads, the number of clean reads in the two libraries was 40,779,290 and 41,776,346. The Q20 and Q30 of the two samples were at least 97.28 and 93.20%, respectively, and the GC contents both exceeded 45%. Additionally, 73.79% or 71.67% of reads of each samples were mapped to the *Populus trichocarpa* Torr. & Gray genome sequence and approximately 47% of the mapped reads were uniquely mapped reads (Table 1).

**Table 1.**
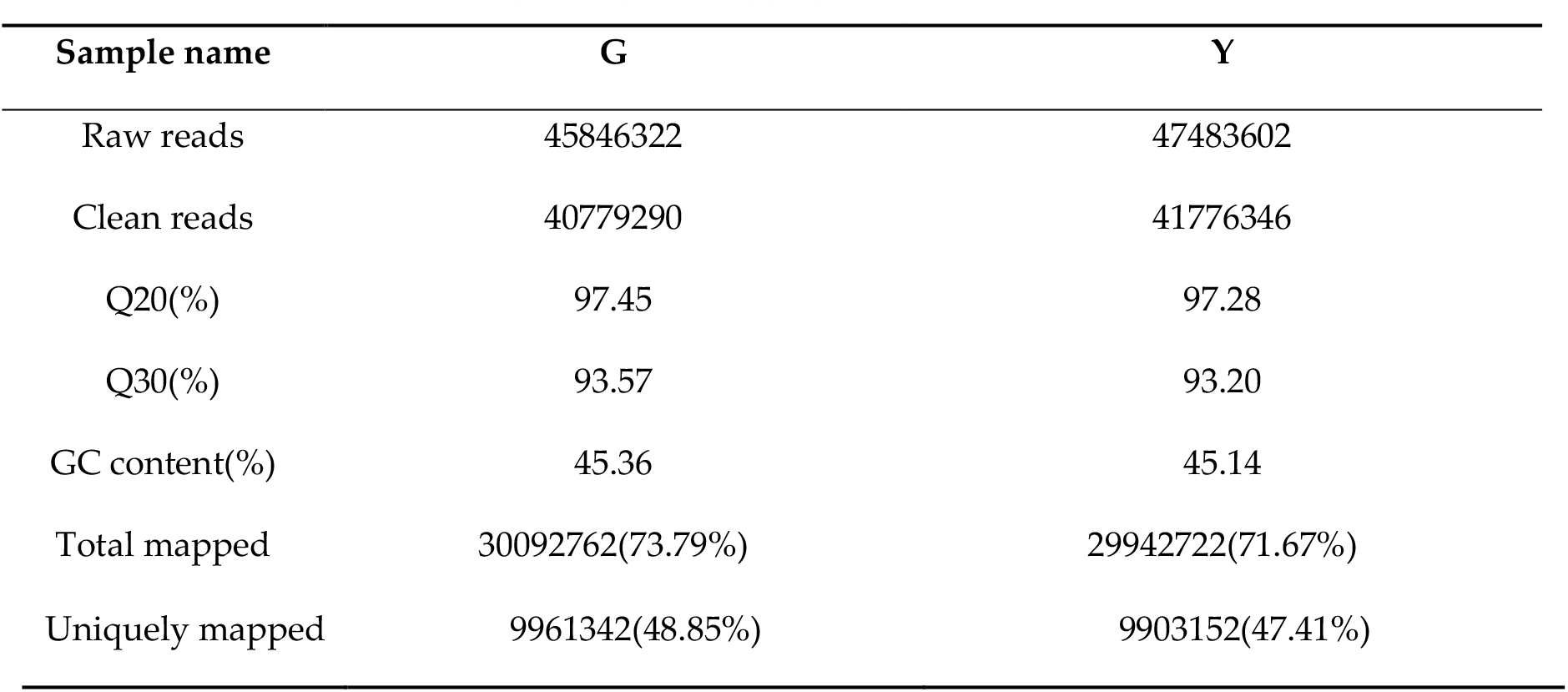
Summary of the sequencing and mapping results.

### 2.2. Analysis of gene expression

In total, the number of expressed genes were 28,657 and 28,124 in green (G) and yellow (Y) leaves, respectively, of which 1760 and 1227 genes were expressed specifically in the G and Y type (Figure 3). In order to identify DEGs between G and Y, we set the expression of genes in G as the control and identified genes that were up-or downregulated in Y. Accordingly, A total of 153 DEGs were found in Y, including 52 up-regulated genes and 101 down-regulated genes.

**Figure 3.**
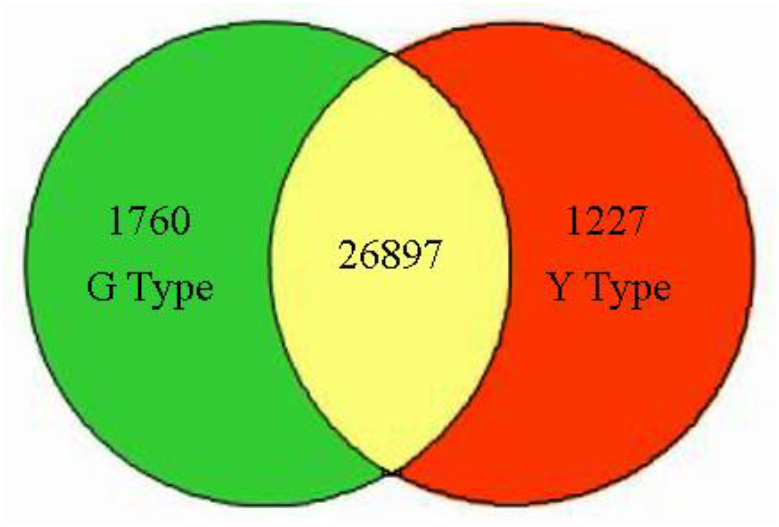
The numbers of specific genes and shared genes between G and Y.

### 2.4. Gene functional annotation by GO, and KEGG

GO assignments were used to classify the functions of DEGs. A total of 12, 9, and 5 of the DEGs were divided into biological processes, cellular components and molecular functions respectively, and some DEGs were annotated with more than one GO term (Figure 4). In the biological process category, a large number of DEGs fell into the categories of ‘cellular process’, ‘metabolic process’, and ‘single-organism process’ (Table S1). The most enriched terms of the cellular component were involved in ‘cell’, ‘cell part’, and ‘membrane’, ‘membrane part’ were also significantly enriched terms (Table S1). Meanwhile, the dominant categories with respect to molecular function group were ‘binding’ and ‘catalytic activity’ (Table S1).

**Figure 4.**
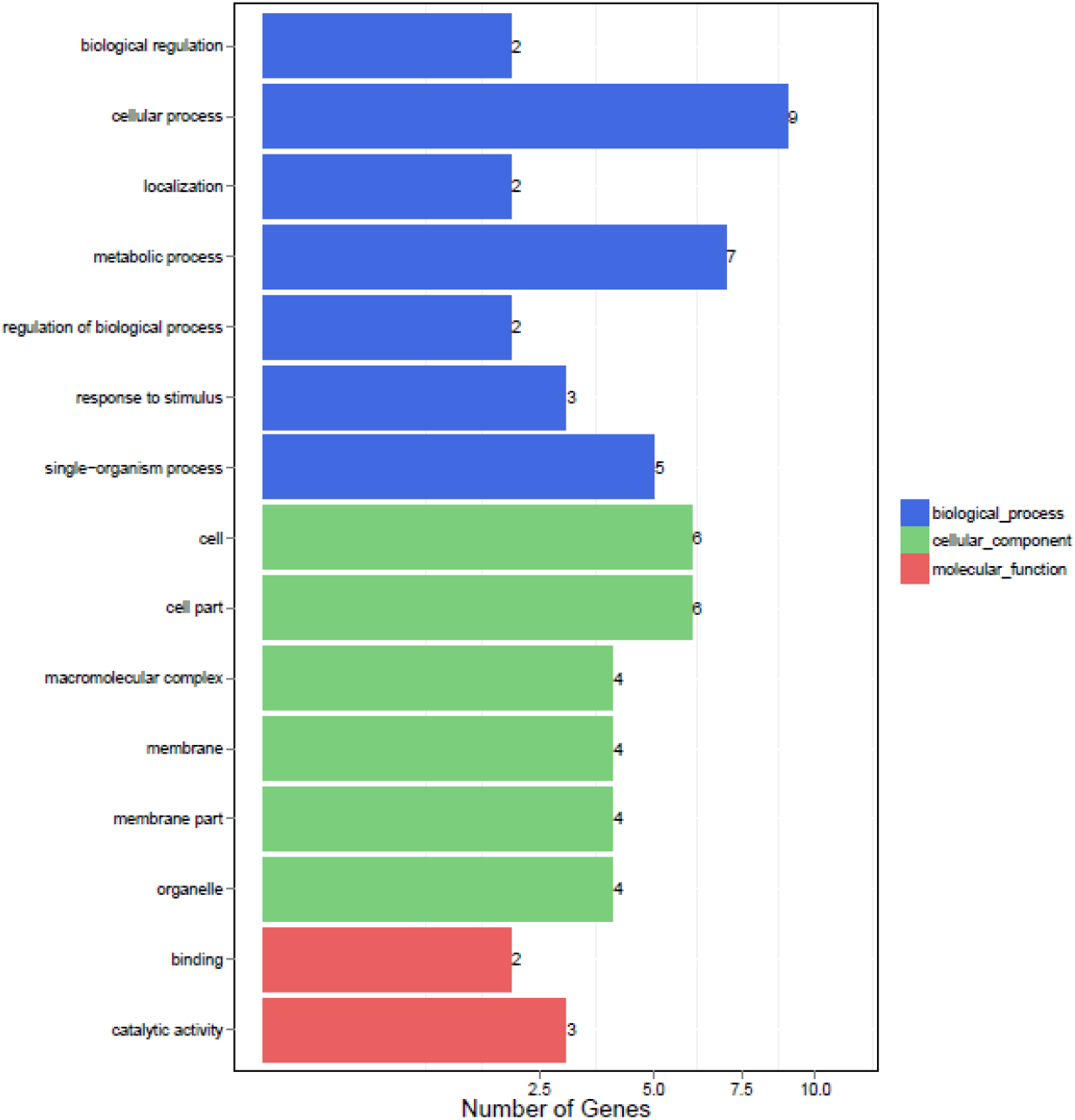
The most enriched GO term assignment to DEGs in G and Y. X-axis displays the gene number. The DEGs were annotated in three categories: biological process, cellular component and molecular function (Y-axis).

KEGG pathway analysis was performed to categorize gene functions with an emphasis on biochemical pathways that were active in G and Y. A total of 52 genes were annotated and assigned to 31 KEGG pathways (Table S2). The most significantly enriched pathway was ‘Metabolic pathways’ (Figure 5), with 15 associated DEGs (ranked by padj value), followed by ‘Biosynthesis of secondary metabolites’ and ‘Ribosome’ with 8 and 5 DEGs, respectively, which supported the results of GO assignments that ‘metabolic process’ was significantly enriched. Moreover, 3 DEGs were assigned to ‘Porphyrin and Chl metabolism’ and 2 DEGs were assigned to ‘Flavonoid biosynthesis’. This cluster of results indicated that the differences in metabolic activities were the main difference between G and Y, and they may perform important roles in the regulating of leaf coloration.

**Figure 5.**
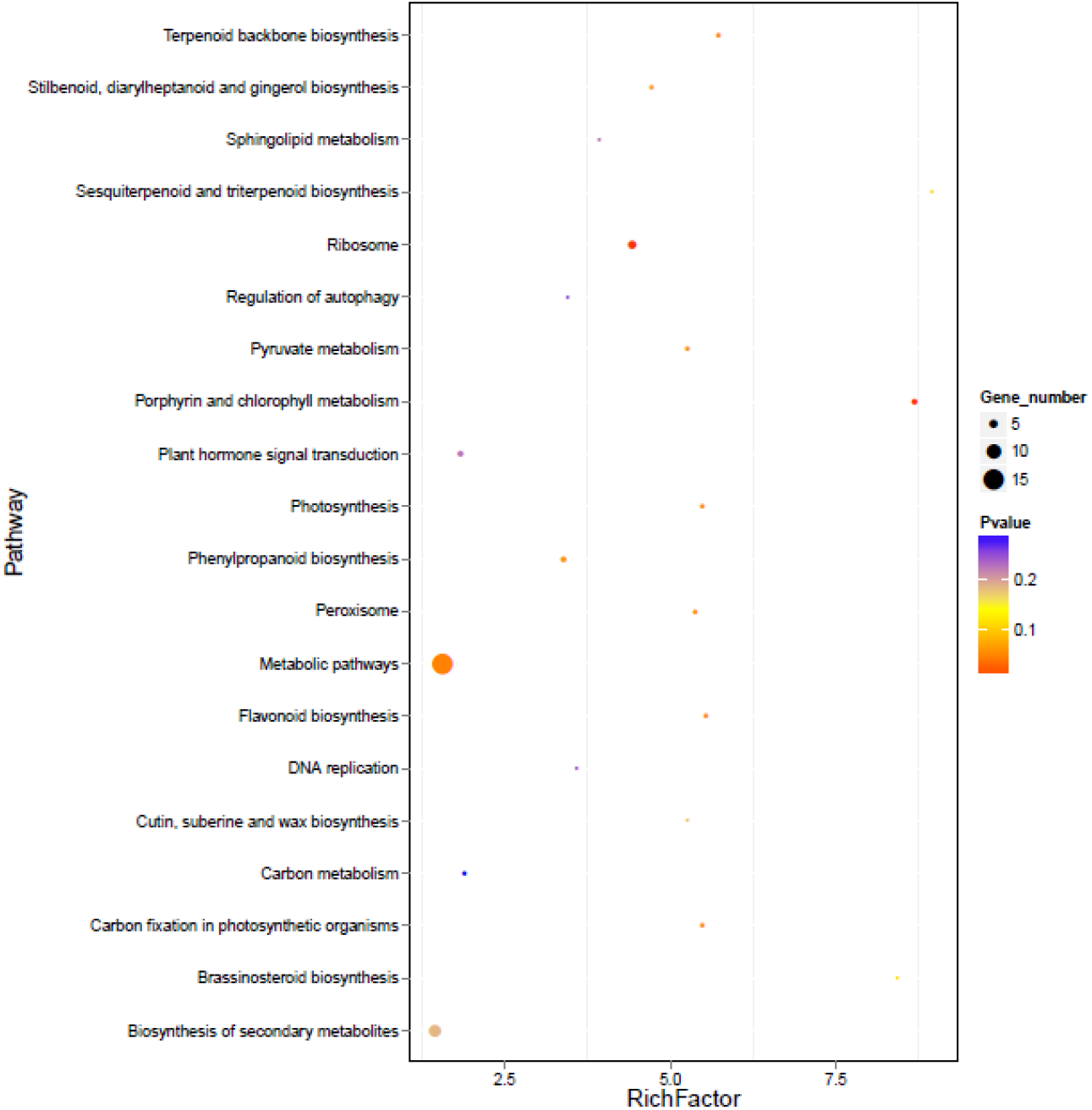
The enriched KEGG pathways of DEGs. The X-axis displays the Rich factor, and the Y-axis displays the KEGG pathways. The size of the dot corresponds to the number of DEGs in the pathway, and the colour of the dot indicates different Q value.

### 2.5. Analysis on Genes Related to Chl and Flavonoid Biosynthesis

Based on the above annotations, we found that the Populus deltoides Marsh transcriptome database contains genes involving in Chl biosynthesis and flavonoid biosynthesis (Table 2). Two genes annotated as CHLP (Potri.019G009000 and Potri.019G024600) were down-regulated in Y. In the last step of Chl a biosynthesis, the geranylgeranyl diphosphate (CHLP, EC:1.3.1.111) catalyzes the reduction of geranylgeranyl pyrophosphate to phytyl pyrophosphate and yields Chl (Figure 6). Furthermore, the gene encoding Chlase (CLH, EC:3.1.1.14) plays roles in the transition of Chl a(b) to Chlide a(b), which was found to be up-regulated in Y. In flavonoid biosynthesis, two genes annotated as HCT were differentially expressed in G and Y. Of these, one gene (Potri.006G034100) was more highly expressed in G while the other gene (Potri.005G028500) was more highly expressed in Y.

**Table 2.**
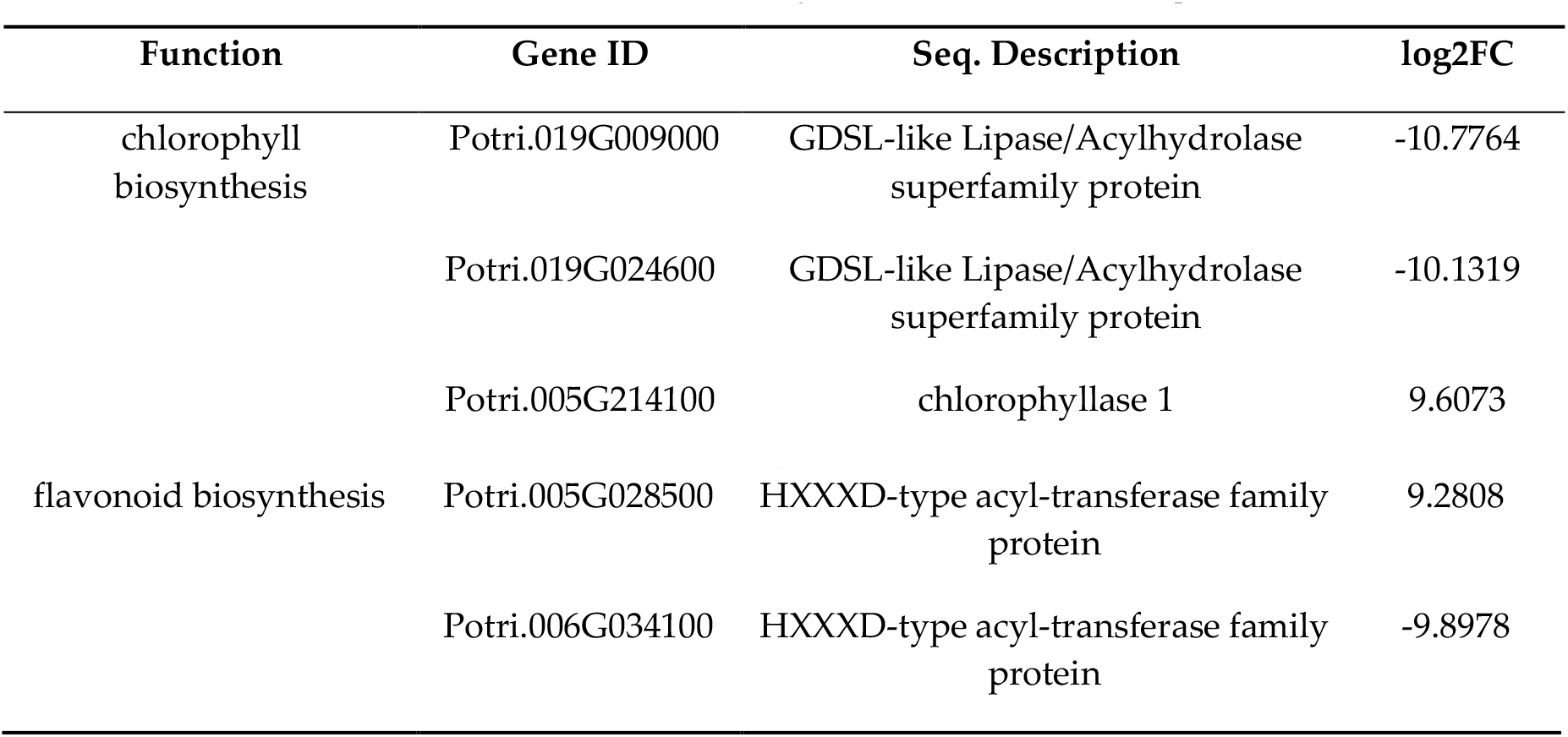
DEGs involved in Chl and flavonoid biosynthesis in mutant transcriptome.

| Function | Gene ID | Seq. Description | log2FC |
| --- | --- | --- | --- |
| chlorophyll biosynthesis | Potri.019G009000 | GDSL-like Lipase/Acylhydrolase superfamily protein | -10.7764 |
|  | Potri.019G024600 | GDSL-like Lipase/Acylhydrolase superfamily protein | -10.1319 |
|  | Potri.005G214100 | chlorophyllase 1 | 9.6073 |
| flavonoid biosynthesis | Potri.005G028500 | HXXXD-type acyl-transferase family protein | 9.2808 |
|  | Potri.006G034100 | HXXXD-type acyl-transferase family protein | -9.8978 |

**Figure 6.**
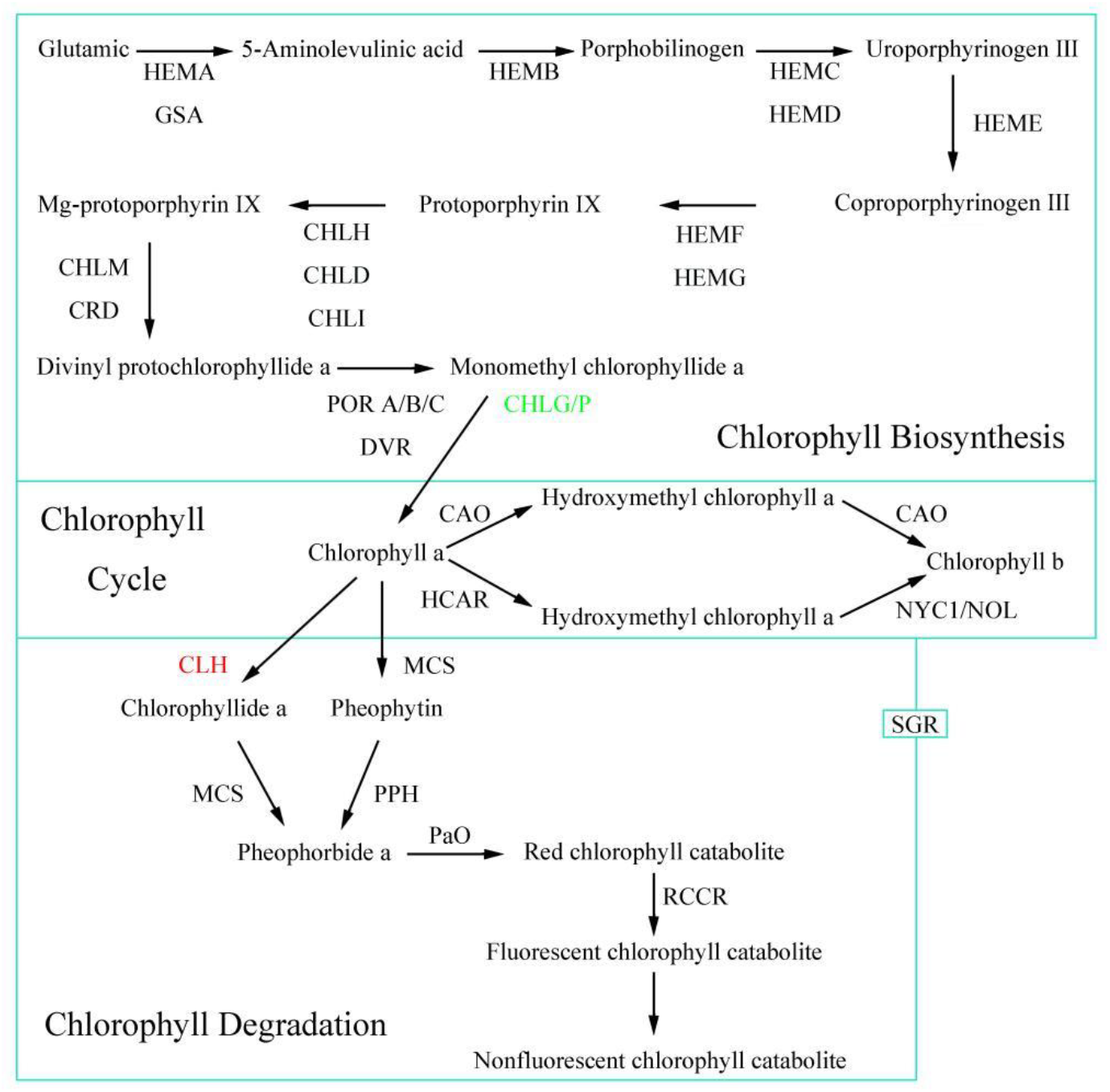
DEGs at the transcript level involved in Chl biosynthesis pathways.Up-regulated genes are marked by red and down-regulated genes by green.

### 2.6. Quantitative real-time PCR validation of RNA Sequencing data

To validate the accuracy of RNA-seq expression results, 8 DEGs with marked changes in plant hormone signal transduction, flavonoid biosynthesis and Chl biosynthesis were detected by qPCR (Figure 7). The results showed that except 3 genes (DELLA, HCT, CLH), the remaining 5 genes were all down-regulated in mutant plants. In general, qRT-PCR results concur with the RNA-seq data, indicating that the DEGs identified by RNA-seq were accurate.

**Figure 7.**
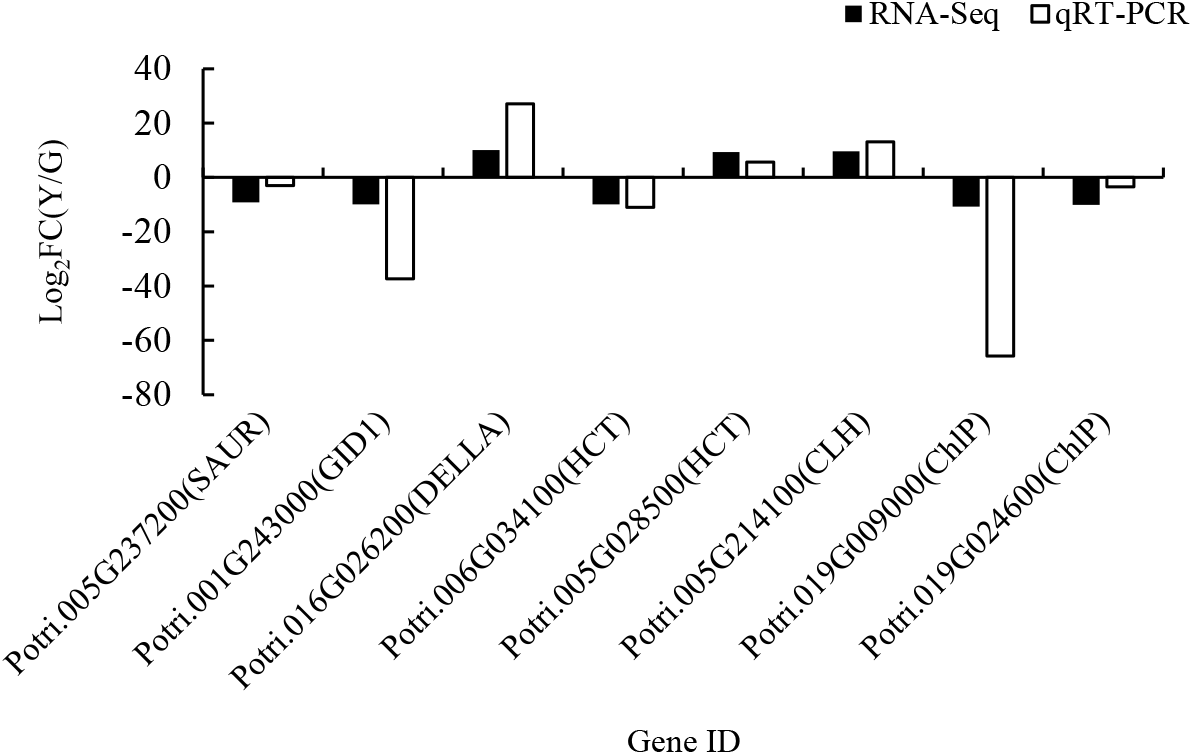
RT-qPCR analysis of the expression values of the 8 DEGs in the Y type. Log2(FC) represents the fold change in Y relative to that in G.

## 3. Discussion

The expression of leaf color in the mutants is often influenced by genes involved in the chloroplast development, Chl synthesis and catabolism, or environmental conditions like temperature and light intensity. Of the many rice yellow green leaf mutants, ygl1 mutant is due to a missense mutation in a highly conserved residue of *YGL1* which encodes the Chl synthase (CHLG) [7], ygl2 mutant is due to an insert mutation of *YELLOW-GREEN LEAF2* which encodes Heme Oxygenase 1 [26]. The impaired chloroplast development of pak-choi yellow leaf mutant is associated with blocked Chl biosynthesis process [21]. In *Setaria italica*, the chlorotic organs is caused by *EGY1* (ethylene-dependent gravitropism-deficient and yellow-green 1), which results in premature senescence and damaged PS II function [27]. The single incompletely dominant gene *Y1718* that is on chromosome 2BS is responsible for the yellow leaf color phenotype of wheat mutant [28]. As a result of a single nucleotide substitution in the *CsChlI* gene for magnesium chelatase I subunit, the cucumber mutant exhibited the golden yellow leaf color throughout its growth stage [25]. The leaf color in the *Japonica* rice is temperature dependent, the mutant displayed yellow-green leaves at low temperature (20°C) and green leaves at higher temperature (34°C) during the seedling stage [29]. The golden leaves of tropical plant *Ficus microcarpa* L. f. cv. is high-light sensitive, which sun-leaves are yellow and shade-leaves are green [30]. In this study, yellow leaf color of G-type is caused by genes involved in the Chl synthesis and catabolism.

### 3.1. The Expression Level of Genes Involved in Chl Biosynthesis Were Altered in Leaf Color Mutants

Leaf color formation is closely related to Chl biosynthesis and breakdown, most leaf color mutations are Chl-deficiency mutations [31]. Chl is responsible for harvesting solar energy and electron transport, even turning plants green because it is Mg^2+^-containing tetrapyrrole pigments [32]. In this study, the novel Chl-deficient chlorina mutant of *Populus deltoides* Marsh with yellow leaf phenotype was identified. Compared with G, the content of photosynthetic pigments in the Y were significantly lower. In particular, the Chl b content were six times higher in Y than G plants. Those results suggesting that the yellow leaf phenotype in mutant is a result of a lack of Chls.

The Chl metabolic process can be subdivided into three parts: biosynthesis of Chl a, the Chl cycle between Chl a and b, and degradation of Chl a [33-35]. Chl is composed of two moieties, Chlide and phytol, which are respectively formed from the precursor molecules 5-aminolevulinate and isopentenyl diphosphate [36]. *CHLP* encodes the enzyme geranylgeranyl reductase catalysing terminal hydrogenation of geranylgeraniol to phytol for Chl synthesis [37,38]. Previous studies revealed that in transgenic tobacco (*Nicotiana tabacum*) expressing antisense *CHLP* RNA, transformants with gradually reduced *CHLP* expression displayed a uniform low pigmentation and a pale or variegated phenotype [39]. In cyanobacterium *Synechocystis* sp. PCC 6803, *∆chlP* mutant exhibit decreased Chl and total carotenoids contents, and unstable photosystems I and II [40]. Two *CHLP* genes (Potri.019G009000 and Potri.019G024600) were identified in our database, and both were down-regulated in the mutant. In the meantime, qPCR experiment further verified that expression levels of *CHLP* genes in Y were highly reduced compared with those in G, which suggesting that a later stage of Chl biosynthesis was interrupted. Parallel experiments also showed that the content of Chlide a was about 4.83% lower, while the content of Chl a was 72.41% lower in the Y compared to G. The result suggests that the inhibition of enzyme activity of CHLP protein is likely to further suppress the biosynthesis of Chl in G. In addition, Our physiological results show that the content of Urogen III in the G is about 4 times than that of the Y, but the content of Coprogen III is no significantly differed between G and Y. Therefore, there might be an interruption between Urogen IIIand Coprogen III during Chl biosynthesis. However, the results need further verification.

Four enzymatic steps of the Chl catabolic pathway are that phytol, magnesium, and the primary cleavage product of the porphyrin ring are catalyzed by Chlase, Mg-dechelatase, pheophorbide a oxygenase, and red Chl catabolite reductase [41]. Chlase catalyzes the hydrolysis ester bond of Chl to yield Chlide and phytol, is thought to be the first enzyme in the Chl degradation [42]. Chlase activity is negatively correlated with Chl levels during citrus fruit color break and Chlase participate in Chl breakdown of citrus [15]. However, some evidence does not support that Chlase play a critical role in Chl degradation during leaf senescence [43-45]. For example, overexpression of *ATHCOR1* which has Chlase activity in *Arabidopsis* leaded to an increased breakdown of Chl a, but the total Chl level was not increased [43]. Similarly, *Arabidopsis* Chlases (*AtCLH1* and *AtCLH2*) is not positively regulated with leaf senescence, *CHL1* and *CHL2* single and double knockout mutant plants do not display a significant delay in senescence [44]. Schelbert et al. also support the opinions that Chlase was not to be essential for dephytylation after Chl is converted into pheophorbide [12]. In our study, the transcript expression patterns suggested that the expression of *CLH* was higher in the Y than in the G. Moreover, previous studies in common wheat (*Triticum aestivum* L.) showed that the gene encoding Chlase in the Chl biosynthesis pathway was also significantly up-regulated in the yellow leaf mutant [45]. Therefore, experiments related to cloning and functional verification of *CLH* in *Populus deltoides* Marsh are need to further verify the function of Chlase in Chl breakdown.

### 3.2. The Expression Level of Genes Involved in Flavonoid Biosynthesis Were Altered in Leaf Color Mutants

Flavonoids, carotenoids, and Chls are the main pigments responsible for flower and leaf color. Previous studies have demonstrated that flavonoids are the main pigments, producing purple, blue, yellow, and red colors in plants [46]. Flavonoids have been known as UV-protecting pigments and antioxidants by scavenging molecular species of active oxygen [47,48]. In *Ficus microcarpa* L. f., the golden leaf mutant is the result of continuous high-light irradiation, and the flavonoid level of golden leaf was 5-fold higher than that of green leaf, the results suggest that the increase of flavonoids in the golden leaf may protect the leaves from high-light stress [49]. In this study, there is no significant differences in the content of flavonoid between Y and G. Therefore, we consider yellow leaf phenotype is caused by genetic factors, not environmental factors. Shikimate/quinate hydroxycinnamoyltransferase (E2.3.1.133, HCT) belongs to the large family of BAHD-like acyltransferases [50]. It is a key enzyme that determines whether 4-coumaroyl CoA is the direct precursor for flavonoid or H-lignin biosynthesis [51]. In Arabidopsis, silencing of the *HCT* gene resulted in severely reduced growth and absent S lignin [52]. The down-regulation of *HCT* have a dramatic effect on lignin content and composition in alfalfa and poplar [53,54]. Up to now, Most studies focus on the effects of *HCT* on lignin synthesis [55,56], while only a few studies related to the *HCT* in flower color or leaf color of plants. It is further proved that the blocked Chl synthesis pathway in Y may be the consequence of yellowing of the leaves.

## 4. Materials and Methods

### 4.1. Plant materials and growing conditions

The green leaf populus cultivar (wild-type) and the yellow leaf populus cultivar (mutant) were used in this study. The plants were three-years-old and grown in Hongxia Nursery, Mianzhu Town, Sichuan Province, China. Leaf tissues were collected in May, sampling three leaves per plant for five plants of each type. All of the leaves were frozen immediately in liquid nitrogen after collection and stored at −80 °C for subsequent experiments.

### 4.2. Measurements of Photosynthetic Pigments, Chl Precursors and flavonoid contents

Approximately 0.1 g leaves of the G and Y were selected for Chl and carotenoid measurements. The pigment (Chl a, Chl b, and carotenoid) contents were measured using the method described by Lichtenthaler [57]. Coprogen III was extracted and determined as described by Bogorad [58]. To measure the contents of Proto IX, Mg-Proto IX, Pchlide and Chlide a, leaves were ground into powders with liquid nitrogen and submerged in nine volumes of phosphate-buffered saline (pH 7.4) in an ice bath, then centrifuged (30 min at 8000 rpm). The supernatant was determined using ELISA kit (MEIMIAN, Jiangsu, China) with a Thermo Scientific Multiskan FC (Thermo Fisher Scientific, MA, USA). Flavonoid contents were measured using a UV1901 PCS Double beam UV-VIS Spectrophotometer (Shanghai Yoke Instrument Co., Ltd., Shanghai, China) according to the instructions of Favonoid Plant kit (Suzhou Comin Biotechnology Co., Ltd., Jiangsu, China). Three biological replicates were evaluated for each sample. The data were analyzed using version 17.0 of SPSS software (SPSS Inc., Chicago, IL, USA) with t test, and means were compared at the significance levels of 0.01 and 0.05. The relative values of photosynthetic pigments and Chl precursors in the Y use the value of G as control and calculated as 1.

### 4.3. RNA extraction, quantification and qualification

Total RNA was isolated from the G and Y leaves using CTAB extraction method. RNA concentration and quality were checked using the Agilent 2100 Bioanalyzer (Agilent Technologies, Santa Clara, USA). RNA purity was measured with a Nano Drop 2000 (Thermo Scientific, USA).

### 4.4. Library preparation for transcriptome sequencing

Two RNA samples were treated with DNaseI to remove any remaining DNA, and then the oligo (dT) magnetic beads were used to collect poly A mRNA fraction. After mixing with fragmentation buffer, the resulting mRNA was broken into short RNA inserts of approximately 200 nt. The fragments were used to synthesize the first cDNA strand via random hexamer priming, and the second-strand cDNA was then synthesized using DNA polymerase I and RNase H. The cDNA fragments was purified using magnetic beads and subjected to end-repair before adding a terminal A at the 3ends. Finally, sequencing adaptors were ligated to the short fragments, which were purified and amplified via polymerase chain reaction (PCR). The two libraries were generated and then sequenced on an Illumina HiSeqTM 4000 platform by Chengdu Life Baseline Technology Co., Ltd. (Chengdu, China).

### 4.5. Quality control and reads mapping

The raw reads were edited to remove adapter sequences, low-quality reads, and reads with >10% of Q < 20 bases, and then mapped using HISAT v2.0.0 software (http://ccb.jhu.edu/software/hisat2/downloads/) to the *Populus trichocarpa* Torr. & Gray genome.

### 4.6. Quantification of gene expression level and differential expression analysis

For gene expression analysis, gene abundance was estimated by RSEM v1.2.30 (http://deweylab.github.io/RSEM/) and then normalized with fragments per kilobase of exon per million mapped reads (FPKM) values [59]. To identify genes that were differently expressed between G and Y, the NOIseq v2.16.0 (http://www.bioconductor.org/packages/release/bioc/html/NOISeq.Html) was used in this experiment. Genes with probability >0.8 and |log2 fold change| ≥ 1 were considered as DEGs between samples.

For functional annotation, GO enrichment analysis of DEGs was performed in the GO database (http://www.geneontology.org/) to calculate gene numbers for every term. The hypergeometric test was conducted to find significantly enriched GO terms in the input list of DEGs. KEGG enrichment analysis was implemented using the database resource (http://www.genome.jp/kegg/). The calculation method of KEGG analysis is the same as the GO analysis.

### 4.7. Real-time RT-PCR

For qPCR analysis, total RNA was extracted using RNAprep Pure Plant Kit (Tiangen Biotech Co. Ltd., Beijing, China), approximately 1 μg RNA was reverse transcribed via a TransScript^®^ All-in-One First-Strand cDNA Synthesis SuperMix for qPCR (Tiangen Biotech Co. Ltd., Beijing, China) according to the manufacturer’s instructions. Eight genes were selected for validation using qRT-PCR. Primer sequences were designed using Primer Premier 5.0 software as shown in Table S3. qPCR DNA amplification and analysis were performed using the *TransScript*^®^ Top Green qPCR SuperMix kit (Tiangen Biotech Co. Ltd., Beijing, China) in accordance with the manufacturer’s protocol with an CFX ConnectTM Real-Time System (Bio-Rad, Hercules, CA, USA). The thermal profile was as follows: pre-denaturation at 94 °C for 30 s; 94 °C for 5 s, 60 °C for 30 s, for 40 cycles. The relative expression level of selected genes in G and Y was normalized to CDC2 and ACT expression. Three biological replicates for each of the reactions were performed. The relative expression levels of target genes were estimated using the 2^−ΔΔCt^ method [60].

### 4.8. Data availability

All the clean reads is available at the National Center for Biotechnology Information (NCBI) Short Read Archive (SRA) Sequence Database (accession number SRA740964). Supplemental files available at FigShare. Table S1 contains significantly enriched gene ontologies among downregulated or upregulated genes in Y type compared to G type. Table S2 contains pathway enrichment. Table S3 contains primers for qPCR analysis.

## 5. Conclusions

In this study, physiological characterization and transcriptome sequence analysis showed that there were distinct differences in coloration between green leaves and yellow mutant leaves of *Populus deltoides* Marsh. Transcriptional sequence analysis identified 5 DEGs that participated in porphyrin and Chl metabolism and flavonoid biosynthesis pathways. Furthermore, RT-qPCR verified that those DEGs were expressed differentially in mutant and wild type plants. Down-regulation of *CHLP* and up-regulation of *CLH* might cause the difference of leaves. These results provide an excellent platform for future studies seeking for the molecular mechanisms underlying the yellowing phenotype in *Populus deltoides* Marsh and other closely related species.

## Author Contributions

F.Z. and S.Z. participated in the conceive and design the experiments; F.Z. supervised the experiments; S.Z. performed the most experimental work and image analyses; X.W, J.C. and Q.L. analyzed the transcriptomic data; X.W., Y. Z. and T. L. prepared the figures and tables; S.Z. wrote the paper. All authors read and approved the final manuscript.

## Abbreviations

Chl: Chlorophyll
DEGs: Differentially expressed genes
CHLP: Geranylgeranyl diphosphate
RT-qPCR: Quantitative real-time PCR
CHLD: Magnesium chelatase subunit D
CHLI: Magnesium chelatase subunit I
PORB: Protochlorophyllide oxidoreductase B
PPH: Pheophytinase
Chlase: Chlorophyllase
Urogen III: Uroporphyrinogen III
Coprogen III: Coproporphyrinogen III
Proto IX: Protoporphyrin IX
Mg-Proto IX: Magnesium protoporphyrin IX
Pchlide: Protochlorophyllide
Chlide: Chlorophyllide
G: Green leaves
Y: Yellow leaves
Caro: Carotenoid
CHLG: Chlorophyll synthase
PCR: polymerase chain reaction
NCBI: National Center for Biotechnology Information
SRA: Short Read Archive
FPKM: Fragments per kilobase of exon per million mapped reads

## References

1. Li, Z.; Wang, J.; Zhang, X.; Xu, L. Comparative transcriptome analysis of anthurium “albama” and its anthocyanin-loss mutant. Plos One 2015, 10, e0119027.

2. Li, W.; Yang, S.; Lu, Z.; He, Z.; Ye, Y.; Zhao, B.; Wang, L.; Jin, B. Cytological, physiological, and transcriptomic analyses of golden leaf coloration in *Ginkgo biloba* L. Hortic. Res. 2018, 5.

3. Tanaka, Y., Sasaki, N., Ohmiya, A. Biosynthesis of plant pigments: anthocyanins, betalains and carotenoids. Plant Journal 2008, 54, 733–749.

4. Tanaka, Y.; Brugliera, F.; Kalc, G.; Senior, M.; Dyson, B.; Nakamura, N.; Katsumoto, Y.; Chandler, S. Flower color modification by engineering of the flavonoid biosynthetic pathway: practical perspectives. Biosci. Biotechnol. Biochem. 2010, 74, 1760–1769.

5. Nagata, N.; Tanaka, R.; Satoh, S.; Tanaka, A. Identification of a vinyl reductase gene for chlorophyll synthesis in *Arabidopsis thaliana* and implications for the evolution of Prochlorococcus species. Plant Cell 2005, 17, 233–240.

6. Li, W.; Tang, S.; Zhang, S.; Shan, J.; Tang, C.; Chen, Q.; Jia, G.; Han, Y.; Zhi, H.; Diao, X. Gene mapping and functional analysis of the novel leaf color gene SiYGL1 in foxtail millet [*Setaria italica* (L.) P. Beauv]. Physiol. Plant 2015, 157, 24–37.

7. Wu, Z.; Zhang, X.; He, B.; Diao, L.; Sheng, S.; Wang, J.; Guo, X.; Su, N.; Wang, L.; Jiang, L.; et al. A chlorophyll-deficient rice mutant with impaired chlorophyllide esterification in chlorophyll biosynthesis. Plant Physiol. 2007, 145, 29–40.

8. Zhu, X.; Shuang, G.; Zhongwei, W.; Qing, D.; Yadi, X.; Tianquan, Z.; Wenqiang, S.; Xianchun, S.; Yinghua, L.; Guanghua, H. Map-based cloning and functional analysis of YGL8, which controls leaf colour in rice (*Oryza sativa* L.). BMC Plant Biol. 2016, 16, 134.

9. Luo, T.; Luo, S.; Araujo, W.L.; Schlicke, H.; Rothbart, M.; Yu, J.; Fan, T.; Fernie, A.R.; Grimm, B.; Luo, M. Virus-induced gene silencing of pea CHLI and CHLD affects tetrapyrrole biosynthesis, chloroplast development and the primary metabolic network. Plant Physiol. Biochem. 2013, 65, 17–26.

10. Yang, Y.L.; Xu, J.; Rao, Y.C.; Zeng, Y.J.; Liu, H.J.; Zheng, T.T.; Zhang, G.-H.; Hu, J.; Guo, L.B.; Qian, Q.; et al. Cloning and functional analysis of pale-green leaf (PGL_10_) in rice (*Oryza sativa* L.). Plant Growth Regul. 2016, 78, 69–77.

11. Hörtensteiner, S. Update on the biochemistry of chlorophyll breakdown. Plant Mol. Biol. 2013, 82, 505–17.

12. Schelbert, S; Aubry, S; Burla, B; Agne, B; Kessler, F; Krupinska K, Hörtensteiner, S. Pheophytin pheophorbide hydrolase (pheophytinase) is involved in chlorophyll breakdown during leaf senescence in Arabidopsis. Plant Cell 2009, 21, 767–85.

13. Kariola, T.; Brader, G.; Li, J.; Palva, E.T. Chlorophyllase 1, a damage control enzyme, affects the balance between defense pathways in plants. Plant Cell 2005, 17, 282–294.

14. Jacob-Wilk, D.; Goldschmidt, D.H.E.E.; Riov, J.; Eyal, Y. Chlorophyll breakdown by chlorophyllase: Isolation and functional expression of the Chlase1 gene from ethylene-treated Citrus fruit and its regulation during development. Plant J. 1999, 20, 653–661.

15. Azoulay Shemer, T.; Harpaz-Saad, S.; Belausov, E.; Lovat, N.; Krokhin, O.; Spicer, V.; Standing, K.G.; Goldschmidt, E.E.; Eyal, Y. Citrus chlorophyllase dynamics at ethylene-induced fruit color-break: A study of chlorophyllase expression, posttranslational processing kinetics, and in situ intracellular localization. Plant Physiol. 2008, 148, 108–118.

16. Jung, K.-H.; Hur, J.; Ryu, C.-H.; Choi, Y.; Chung, Y.-Y.; Miyao, A.; Hirochika, H.; An, G. Characterization of a Rice Chlorophyll-Deficient Mutant Using the T-DNA Gene-Trap System. Plant Cell Physiol. 2003, 44, 463–472.

17. Wang, P.; Gao, J.; Wan, C.; Zhang, F.; Xu, Z.; Huang, X.; Sun, X; Deng, X. Divinyl chlorophyll(ide) a can be converted to monovinyl chlorophyll(ide) a by a divinyl reductase in rice. Plant Physio. 2010, 153, 994–1003.

18. Liu, W.; Fu, Y.; Hu, G.; Si, H.; Zhu, L.; Wu, C.; Sun, Z. Identification and fine mapping of a thermo-sensitive chlorophyll deficient mutant in rice (*oryza sativa* L.). Planta 2007, 226, 785–795.

19. Sakuraba, Y.; Rahman, M.L.; Cho, S.H.; Kim, Y.S.; Koh, H.J.; Yoo, S.C.; Paek N.C. The rice faded green leaf locus encodes protochlorophyllide oxidoreductaseb and is essential for chlorophyll synthesis under high light conditions. Plant J. 2013, 74, 122–133.

20. Sakuraba, Y.; Park, S.Y.; Kim, Y.S.; Wang, S.H.; Yoo, S.C.; Hrtensteiner, S.; Paek, N.C. Arabidopsis stay-green2 is a negative regulator of chlorophyll degradation during leaf senescence. Mol. Plant. 2014, 7, 1288–1302.

21. Zhang, K.; Liu, Z.; Shan, X.; Li, C.; Tang, X.; Chi, M.; Feng, H. Physiological properties and chlorophyll biosynthesis in a pak-choi (*Brassica rapa* L. *ssp*. *chinensis*) yellow leaf mutant, pylm. Acta Physiol. Plant. 2017, 39, 22.

22. Akifumi, A.; Shozo, K.; Nami, G.Y.; Mikio, S.; Nobuhito, M.; Hiroshi, Y.; Yoshiko, K. Color recovery in berries of grape (*vitis vinifera* L.) ‘benitaka’, a bud sport of ‘italia’, is caused by a novel allele at the *VvmybA1* locus. Plant Sci. 2009, 176, 470–478.

23. Wasscher, J. The importance of sports in some florist’s flowers. Euphytica 1956, 5, 163–170.

24. Li, C.F.; Xu, Y.X.; Ma, J.Q.; Jin, J.Q.; Huang, D.J.; Yao, M.Z.; Ma, C.L.; Chen, L. Biochemical and transcriptomic analyses reveal different metabolite biosynthesis profiles among three color and developmental stages in ‘Anji Baicha’ (*camellia sinensis*). BMC Plant Bio. 2016, 16, 195.

25. Gao, M.; Hu, L.; Li, Y.; Weng, Y. The chlorophyll-deficient golden leaf mutation in cucumber is due to a single nucleotide substitution in *CsChlI* for magnesium chelatase I subunit. Theor. Appl. Genet. 2016, 129, 1–13.

26. Chen, H.; Cheng, Z.; Ma, X.; Wu, H.; Liu, Y.; Zhou, K.; Chen, Y.; Ma, W.; Bi, J.; Zhang, X.; et al. A knockdown mutation of *YELLOW-GREEN LEAF2* blocks chlorophyll biosynthesis in rice. Plant Cell Rep. 2013, 32, 1855–1867.

27. Zhang, S.; Zhi, H.; Li, W.; Shan, J.; Tang, C.; Jia, G.; Tang, S.; Diao, X. SiYGL2 is involved in the regulation of leaf senescence and photosystem II efficiency in *Setaria italica* (L.) P. Beauv. Front. Plant Sci. 2018, 9, 1308.

28. Zhang, L.; Liu, C.; An, X.; Wu, H.; Feng, Y.; Wang, H.; Sun, D. Identification and genetic mapping of a novel incompletely dominant yellow leaf color gene, *Y1718*, on chromosome 2bs in wheat. Euphytica 2017, 213, 141.

29. Ruan, B.; Gao, Z.; Zhao, J.; Zhang, B.; Zhang, A.; Hong, K.; Yang, S.; Jiang, H.; Liu, C.; Chen, G.; et al. The rice *YCL* gene encoding an Mg2+-chelatase ChlD subunit is affected by temperature for chlorophyll biosynthesis. J. Plant Biol. 2017, 60, 314–321.

30. Yoshinobu, K.; Yasusi, Y.; Yasuko, S.; Ayumu, T.; Shunichi, T.; Hideo, Y. High-susceptibility of photosynthesis to photoinhibition in the tropical plant *Ficus microcarpa* L. f. cv. golden leaves. BMC Plant Biol. 2002, 2, 2.

31. Wang, L.; Yue, C.; Cao, H.; Zhou, Y.; Zeng, J.; Yang, Y.; Wang, X. Biochemical and transcriptome analyses of a novel chlorophyll-deficient chlorina tea plant cultivar. BMC Plant Physiol. 2014, 14, 352.

32. Tanaka, A.; Tanaka, R. Chlorophyll metabolism. Curr. Opin. Plant Biol. 2006, 9, 248–255.

33. Eckhardt, U.; Grimm, B.; Hortensteiner, S. Recent advances in chlorophyll biosynthesis and breakdown in higher plants. Plant Mol Biol. 2004, 56, 1–14.

34. Tanaka, A.; Ito, H.; Tanaka, R.; Tanaka, N.K.; Yoshida, K.; Okada, K. Chlorophyll a oxygenase (cao) is involved in chlorophyll b formation from chlorophyll a. Proc. Natl. Acad. Sci. 1998, 95, 12719–12723.

35. Masuda, T.; Fujita, Y. Regulation and evolution of chlorophyll metabolism. Photochem Photobiol Sci. 2008, 7, 1131–1149.

36. von Wettstein, D.; Gough, S.; Kannangara, C.G. Chlorophyll biosynthesis. Plant Cell 1995, 7, 1039–1057.

37. Addlesee, H.A.; Gibson, L.C.; Jensen, P.E.; Hunter, C.N. Cloning, sequencing and functional assignment of the chlorophyll biosynthesis gene, chlp, of *Synechocystis* sp. pcc 6803. FEBS Lett. 1996, 389, 126–130.

38. Addlesee, H.A.; Hunter, C.N. Physical mapping and functional assignment of the geranylgeranyl-bacteriochlorophyll reductase gene, bchp, of rhodobacter sphaeroides. J. Bacteriol. 1999, 181, 7248.

39. Tanaka, R.; Oster, U.; Kruse, E.; Rudiger, W.; Grimm, B. Reduced activity of geranylgeranyl reductase leads to loss of chlorophyll and tocopherol and to partially geranylgeranylated chlorophyll in transgenic tobacco plants expressing antisense rna for geranylgeranyl reductase. Plant Physiol. 1999, 120, 695–704.

40. Shpilyov, A.V.; Zinchenko, V.V.; Shestakov, S.V.; Grimm, B.; Lokstein, H. Inactivation of the geranylgeranyl reductase (chlp) gene in the cyanobacterium *Synechocystis* sp. pcc 6803. BBA - Bioenergetics 2005, 1706, 195–203.

41. Harpazsaad, S.; Azoulay, T.; Arazi, T.; Benyaakov, E.; Mett, A.; Shiboleth, Y.M.; Hortensteiner, S.; Gidoni, D.; Galon, A.;Goldschmidt, E.E.; et al. Chlorophyllase is a rate-limiting enzyme in chlorophyll catabolism and is posttranslationally regulated. Plant Cell 2007, 19, 1007–1022.

42. Tsuchiya, T.; Ohta, H.; Okawa, K.; Iwamatsu, A.; Shimada, H.; Masuda, T.;Takamiya, K. Cloning of chlorophyllase, the key enzyme in chlorophyll degradation: finding of a lipase motif and the induction by methyl jasmonate. Proc. Natl. Acad. Sci. USA 1999, 96, 15362–15367.

43. Benedetti, C.E.; Arruda, P. Altering the expression of the chlorophyllase gene ATHCOR1 in transgenic *Arabidopsis* caused changes in the chlorophyll-to-chlorophyllide ratio. Plant Physiol. 2002, 128, 1255–1263.

44. Schenk, N.; Schelbert, S.; Kanwischer, M.; Goldschmidt, E.E.; Dormann, P.; Hortensteiner, S. The chlorophyllases ATCLH1 and ATCLH2 are not essential for senescence-related chlorophyll breakdown in *Arabidopsis thaliana*. FEBS Lett. 2007, 581, 5517–5525.

45. Wu, H.; Shi, N.; An, X.; Liu, C.; Fu, H.; Cao, L.; Feng, Y.; Sun, D.; Zhang, L. Candidate genes for yellow leaf color in common wheat (*Triticum aestivum* L.) and major related metabolic pathways according to transcriptome profiling. Int. J. Mol. Sci. 2018, 19, 1594.

46. Sobel, J.M.; Streisfeld, M.A. Flower color as a model system for studies of plant evo-devo. Front. Plant Sci. 2013, 4, 321.

47. Schmelzer, E.; Jahnen, W.; Hahlbrock, K. In situ localization of light-induced chalcone synthase mrna, chalcone synthase, and flavonoid end products in epidermal cells of parsley leaves. Proc. Natl. Acad. Sci. USA. 1988, 85, 2989–2993.

48. Hernández; Iker; Alegre; Leonor; Breusegem, V.; Frank. How relevant are flavonoids as antioxidants in plants? Trends Plant Sci. 2009, 14, 125–132.

49. Yamasaki, H., Heshiki, R. N. Leaf-goldenning induced by high light in *Ficus microcarpa* L.f., a tropical fig. J. Plant Res. 1995, 108, 171–180.

50. D’Auria, J.C. Acyltransferases in plants: a good time to be BAHD. Curr. Opin. Plant Biol. 2006, 9, 331–340.

51. Li, X.; Bonawitz, N.D.; Weng, J.K.; Chapple, C. The growth reduction associated with repressed lignin biosynthesis in arabidopsis thaliana is independent of flavonoids. Plant Cell 2010, 22, 1620–1632.

52. Hoffmann, L.; Besseau, S.; Geoffroy, P.; Ritzenthaler, C.; Meyer, D.; Lapierre, C.; Pollet, B.; Legrand, M. Silencing of hydroxycinnamoyl-coenzyme a shikimate/quinate hydroxycinnamoyltransferase affects phenylpropanoid biosynthesis. Plant Cell 2004, 16, 1446–1465.

53. Pu, Y.; Chen, F.; Ziebell, A.; Davison, B.H.; Ragauskas, A.J. NMR characterization of C3H and HCT down-regulated alfalfa lignin. BioEnergy Res. 2009, 2, 198.

54. Peng, X.; Sun, S.; Wen, J.; Yin, W.; Sun, R. Structural characterization of lignins from hydroxycinnamoyl transferase (HCT) down-regulated transgenic poplars. Fuel 2014, 134, 485–492.

55. Cui, K.; Wang, H.; Liao, S.; Tang, Q.; Li, L.; Cui, Y.; He Y. Transcriptome sequencing and analysis for culm elongation of the world’s largest bamboo (*dendrocalamus sinicus*). Plos One 2016, 11, e0157362.

56. Wu, Y.F.; Zhao, Y.; Liu, X.Y.; Gao, S.; Cheng, A.X.; Lou, H.X. Isolation and functional characterization of hydroxycinnamoyltransferases from the liverworts plagiochasma appendiculatum, and marchantia paleacea. Plant Physiol. Biochem. 2018, 129, 400–410.

57. Lichtenthaler, H.K. Chlorophylls and carotenoids: Pigments of photosynthetic biomembranes. Methods Enzymol. 1987, 148, 350–382.

58. Bogorad, L. Porphyrin synthesis. In Methods in Enzymology; Colowick, S.P., Kaplan, N.O., Eds.; Academic Press: New York, NY, USA, 1962; Volume 5, pp. 885–895, ISBN 0076-6879.

59. Mortazavi, A.; Williams, B.A.; McCue, K.; Schaeffer, L.; Wold, B. Mapping and quantifying mammalian transcriptomes by RNA-Seq. Nat Methods 2008, 5, 621–628.

60. Schmittgen, T.D.; Livak, K.J. Analyzing real-time PCR data by the comparative CT method. Nat. Protoc. 2008, 3, 1101–1108.

